# Pollinator niche partitioning and asymmetric facilitation contribute to the maintenance of diversity

**DOI:** 10.1101/2020.03.02.974022

**Authors:** Na Wei, Rainee L. Kaczorowski, Gerardo Arceo-Gómez, Elizabeth M. O’Neill, Rebecca A. Hayes, Tia-Lynn Ashman

**Affiliations:** Department of Biological Sciences, University of Pittsburgh, Pittsburgh, PA, USA.; The Holden Arboretum, Kirtland, OH, USA.; Department of Biological Sciences, East Tennessee State University, Johnson City, TN, USA.

## Abstract

Mechanisms that favor rare species are key to the maintenance of diversity. One of the most critical tasks for biodiversity conservation is understanding how plant–pollinator mutualisms contribute to the persistence of rare species, yet this remains poorly understood. Using a process-based model that integrates plant–pollinator and interspecific pollen transfer networks with floral functional traits, we show that niche partitioning in pollinator use and asymmetric facilitation confer fitness advantage of rare species in a biodiversity hotspot. While co-flowering species filtered pollinators via floral traits, rare species showed greater pollinator specialization leading to higher pollination-mediated male and female fitness than abundant species. When plants shared pollinator resources, asymmetric facilitation via pollen transport dynamics benefited the rare species at the cost of the abundant ones, serving as an alternative diversity-promoting mechanism. Our results emphasize the importance of community-wide plant–pollinator interactions that affect reproduction for biodiversity maintenance.

## Main Text

How numerous rare species coexist with abundant species is a major unresolved question in ecology but is essential to understanding the maintenance of species diversity (*1, 2*). Plant– pollinator interactions are key to the diversification of flowering plants (*3*) and have been identified among the most important drivers of biodiversity on Earth (*4*). Yet we still lack a clear view as to how community-wide interactions between plants and pollinators contribute to the persistence of rare species that are at greater extinction risk than more abundant ones (*5–8*). Mechanisms such as niche partitioning (*9*) and facilitation (*10*) can help maintain rare species (*2*). Niche partitioning can prevent interspecific competitive exclusion between rare and abundant species. Facilitation, on the other hand, generates positive interspecific interactions. Both mechanisms can operate at the pollination stage of the plant life cycle and confer pollination-mediated fitness advantage to the rare species over abundant ones.

Tracking pollination-mediated fitness is, however, more complex than tracking fitness at later life stages (e.g. seed production or seedling growth). Because most plants are hermaphrodite (*11*), fitness at the pollination stage has both female and male components (via ovules that house eggs and pollen that houses sperm). Thus, fitness gain is only achieved from a female–male interaction when pollen from conspecific donors reaches the ovules. The receipt of conspecific pollen (CP) per ovule can therefore reflect this joint fitness gain mediated by pollinators. In contrast, female fitness loss occurs when the pollen received is from another species [i.e. heterospecific pollen (HP) receipt, (*12*)], displacing or interfering with legitimate pollination. Likewise, male fitness loss occurs when pollinators misdeliver pollen, that is, transport it to heterospecific rather than conspecific recipients [i.e. CP misplacement, (*13*)]. Given the multiple pathways of fitness accrual via complex plant–pollinator interactions, a community-wide study is required to assess how these combine to affect diversity maintenance; yet no such study exists.

As pollinator service is often limited in nature (*14, 15*), competition for successful pollination predicts limiting similarity in pollinator sharing (*6*). Such pollinator niche partitioning can potentially favor rare plant species via greater specialization than abundant species, because of fitness costs associated with generalization. In a diverse co-flowering plant community (Fig. 1A), the cost of being a generalist includes high risks of male fitness loss due to CP misplacement to heterospecifics and female fitness loss due to HP receipt, thus reducing joint fitness gain (*12*). In contrast, the benefit of being a specialist may include improved CP delivery by pollinators and lower risks of male and female fitness loss.

**Fig. 1.**
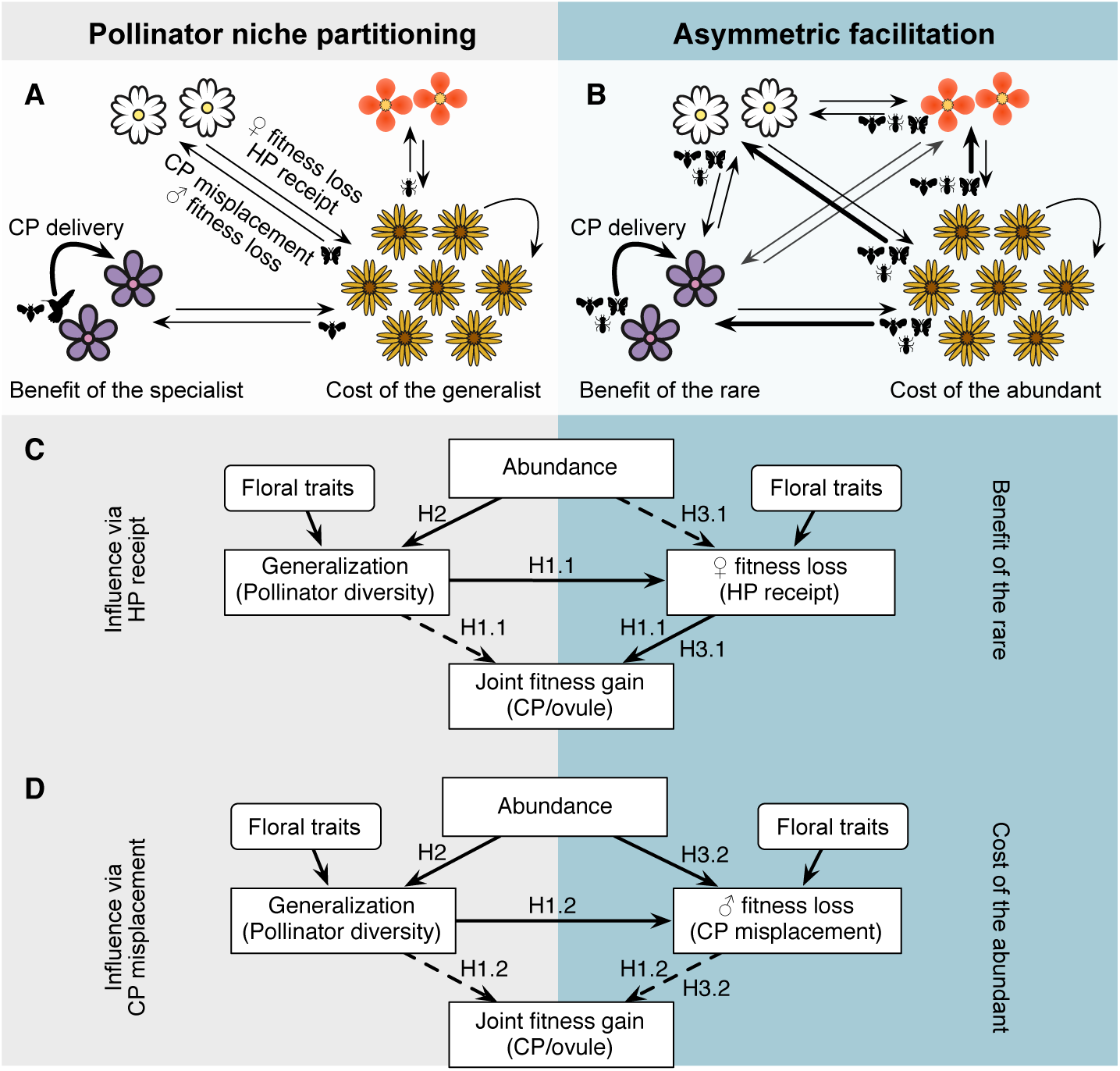
Mechanisms by which pollinator niche partitioning and asymmetric facilitation confer rare species advantage. (**A**) Niche partitioning results from differences in pollinator use by plants. Generalist plants by virtue of sharing pollinators with other species experience high risks of conspecific pollen (CP) misplacement (i.e. male fitness loss) and heterospecific pollen (HP) receipt (i.e. female fitness loss). In contrast, specialist plants benefit from lower risks of fitness losses and higher chances of CP delivery (thickened curved arrow). Niche partitioning can favor rare species via greater specialization in rare species relative to abundant ones. (**B**) When sharing pollinator niche, abundant species experience higher CP misplacement (thickened straight arrow) when pollinators that primarily visit abundant species move to rare species. In turn, rare species benefit from pollinators attracted by abundant species, leading to greater CP receipt than if they were growing alone. (**C** and **D**) The two mutually non-exclusive mechanisms can be examined by linking pollinator niche breath (generalization) and plant rarity (abundance) to HP receipt (**C**) and CP misplacement (**D**), which influence joint fitness gain. Links indicate positive (solid) and negative (dashed) relationships. To support the hypothesis of rare species advantage due to pollinator niche difference, abundance is expected to be positively linked to generalization (H2) and generalization is negatively linked to joint fitness gain (i.e. fitness costs of generalization; H1.1 and H1.2 net effects). To support the hypothesis of asymmetric facilitation, rare species are expected to be facilitated more by receiving CP along HP, leading to increased joint fitness gain (H3.1), and abundance species as facilitators are expected to experience higher male fitness loss, leading to reduced joint fitness gain (H3.2).

When pollinator niches however overlap, asymmetric facilitation can favor rare species (*8, 16*) at the expense of abundant ones (Fig. 1B). Rare species benefit from pollinators attracted by abundant heterospecific neighbors (*8*). Although rare species may also receive HP when sharing pollinators with abundant species (*17*), they will receive more CP than they would if growing alone [i.e. a positive relation between CP–HP receipt; (*16, 18*)]. Abundant species, as facilitators, may experience more CP misplacement to heterospecifics than they would if growing without rare species. Thus, asymmetric facilitation can potentially increase the joint fitness gain of rare species but decrease that of abundant ones due to higher male fitness loss.

Theory predicts that functional trait divergence among species that share pollinator resources leads to increasing diversity (*19*). Traits that filter pollinators by floral advertisement and mechanical fit are important in mediating pollinator niche (*20*). Female and male function traits such as stigma and pollen features can influence pollen receipt and donation, as well as reward signaling (*12, 21*). However, evidence linking these floral functional traits to pollinator niche difference (*22*), male (*23*) and female fitness loss (*18*), and joint fitness gain beyond pairs of interacting species is rare, and virtually nonexistent across entire interaction networks in species-rich communities. This is perhaps due to the challenges of recording pollination-mediated fitness of all the taxa in these communities, especially identifying and tracking misdelivered pollen grains (*13*). Thus, the combinations of traits that govern pollinator diversity and fitness differences among plant species, and thereby modulate the strength of niche partitioning and facilitation, remain entirely unknown in the very communities where we expect these processes to be the strongest – high diversity ecosystems such as global biodiversity hotspots.

Here we evaluated the two mutually non-exclusive mechanisms (i.e. pollinator niche partitioning and asymmetric facilitation) hypothesized to underlie rare species advantage, along with potential functional trait drivers, in the serpentine seeps of California, USA, a global biodiversity hotspot (*24*). We formulated a process-based model (Fig. 1 C and D) to describe the relationships among functional traits and pathways for fitness gains and losses associated with pollinator niche breath (generalization) and plant rarity (abundance) across all the species in the co-flowering community. To assess the hypothesis of rare species advantage due to pollinator niche partitioning, we first tested whether species are limited in pollinator sharing. We then asked whether rare species are more specialized (Fig. 1 C and D, H2) leading to higher joint fitness gain than abundant species (net effects of H1.1 and H1.2), and whether this is the result of (both male and female fitness) costs associated with generalization. To assess the hypothesis of asymmetric facilitation, we asked whether rare species are facilitated more than abundant ones by receiving more CP (along with HP), leading to increased joint fitness gain (Fig. 1C, H3.1). In turn, we asked whether abundant species, as facilitators, experience higher male fitness loss, leading to reduced joint fitness gain relative to rare species (Fig. 1D, H3.2). These hypotheses were examined using phylogenetic structural equation modeling (PSEM) that leveraged species-specific metrics derived from community-wide plant–pollinator and interspecific pollen transfer networks, and a suite of floral functional traits.

We observed plant–pollinator interactions during two consecutive flowering seasons in a system of serpentine seeps (10,000 m^2^) at the McLaughlin Natural Reserve (38.8582 °N, 122.4093 °W; table S1). Among the 7324 pollinators that visited 79 co-flowering plant species (of 62 genera from 29 families) (Fig. 2A), 416 species were identified (table S2): 192 bees (*n* = 4951 individuals; Hymenoptera), 131 flies (*n* = 1409; Diptera), 35 beetles (*n* = 428; Coleoptera), 30 butterflies and moths (*n* = 244; Lepidoptera), 14 wasps (*n* = 104; Hymenoptera), 3 ants (*n* = 22; Hymenoptera), 10 other insect species (*n* = 25), and 1 hummingbird species (*n* = 141; Trochilidae). The plant–pollinator network (Fig. 2A), based on sufficient field observations (fig. S1, rarefaction), revealed substantial variation in pollinator niche breath among plant species, ranging from a plant interacting with one to 77 pollinator species (mean = 23). Different from the nested structure of many ecological networks (*25*) where specialists share and interact with only subsets of the partners of generalists, the network was significantly less nested (*P* < 0.001; table S3), minimizing pollinator sharing. These plants together showed significantly less niche overlap (Horn’s index = 0.086, null mean = 0.412, *P* < 0.001) and fewer shared pollinator partners (observed mean = 3, null mean = 17, *P* < 0.001) than expected by random. For individual plant species, the majority (85%) exhibited significantly higher degrees of specialization than random expectations (e.g. pollinator diversity, *P* < 0.001; dissimilarity between pollinator use and pollinator species pool, *P* < 0.01; table S3), while acknowledging weak differences in species flowering phenology (*26*) that may influence the availability of pollinator partners. Importantly, as hypothesized, rare species were more specialized than abundant ones (PSEM, *r* = 0.41, *P* < 0.05; Fig. 3). Overall, these plant–pollinator interactions strongly demonstrate pollinator niche difference among coexisting plants.

**Fig. 2.**
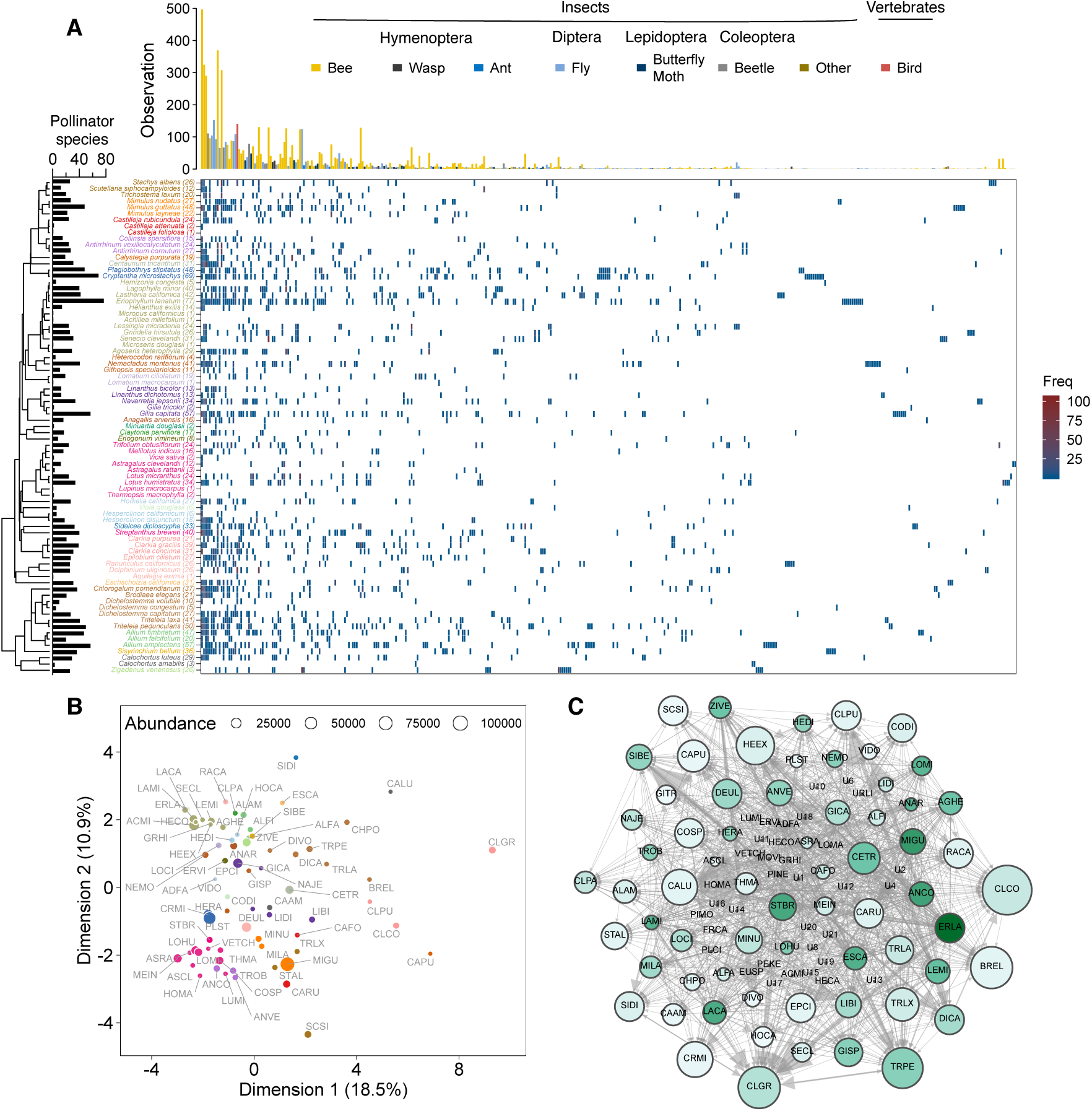
Community-wide plant–pollinator and interspecific pollen transfer networks and floral functional traits. (**A**) The lack of nestedness in the plant–pollinator network provides evidence for pollinator niche partitioning. Plant species (*n* = 79) colored by families are arranged on the left according to phylogeny. The number of pollinator species that each plant interacted with are shown as black bars and numbers within parentheses. Pollinator species (*n* = 416) are arranged along the top according to the size and similarity of plant assemblages that they interacted with. (**B**) Plant species (abbreviated as the first two letters of genus and species names and colored by plant family, *n* = 73) are segregated along the first two dimensions, which represent mainly size-related and other (shape/color/inflorescence) floral traits, respectively, in a multivariate analysis of 20 floral traits (see table S4 for trait details). These traits vary independently from species floral abundance [symbol size, see (*27*)]. (**C**) Pollen transfer network was based on pollen deposited on 54 stigmas of 66 individual plant species. Plant species are nodes in the network. Node size indicates the number of species that pollen is received from, and node color darkness indicates the number of species that pollen is donated to. That is, larger nodes represent better recipients and darker nodes better donors. Arrows and their sizes indicate the direction and amount of pollen transfer, respectively. Species abbreviations follow (B), with those unidentified species denoted with ‘U’.

**Fig. 3.**
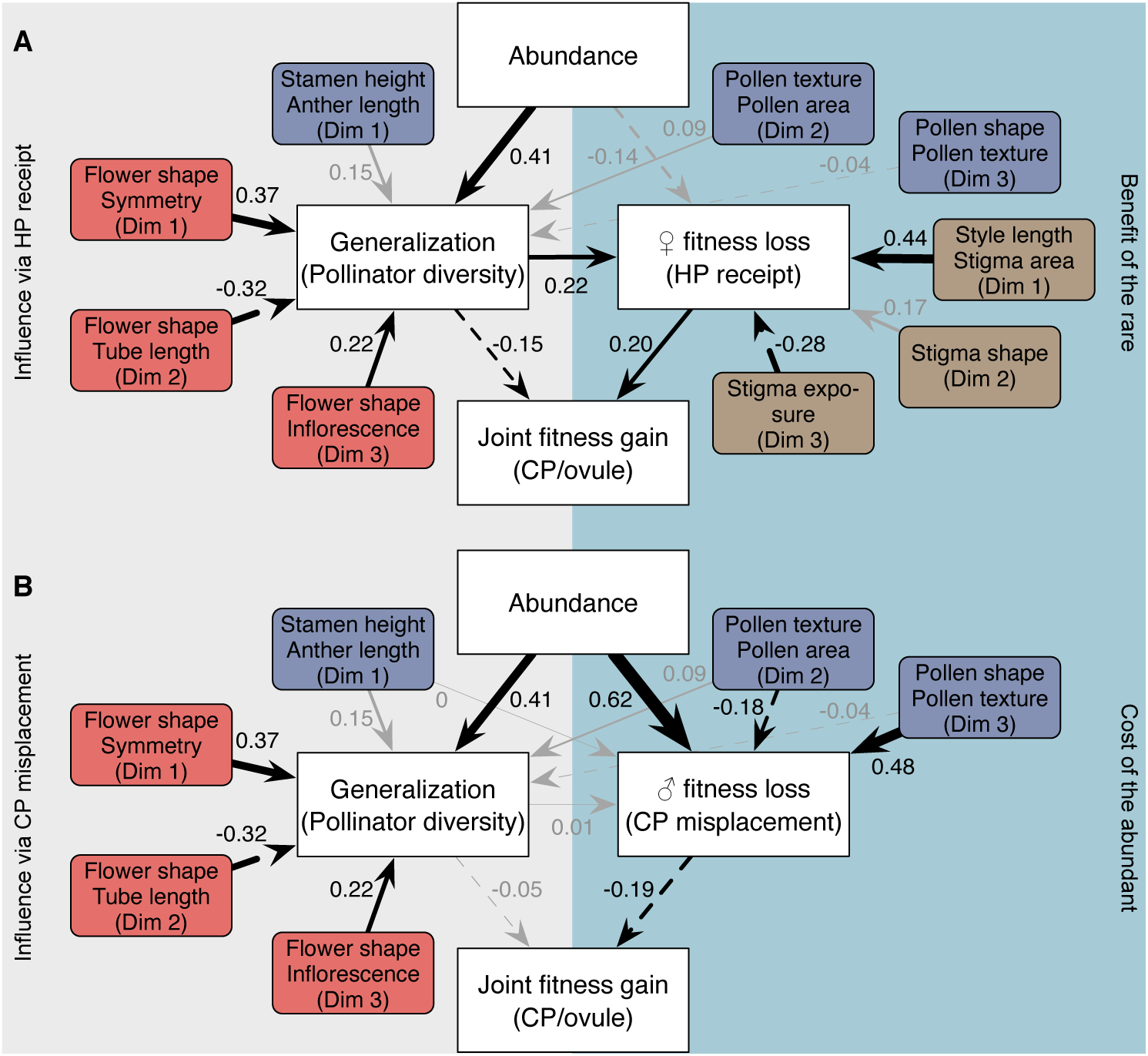
Plant–pollinator interactions favor rare species through pollinator niche partitioning and asymmetric facilitation which are mediated by floral functional traits. Results of phylogenetic structural equation modeling demonstrate how pollinator niche breath (generalization) and plant rarity (abundance) affect fitness loss via heterospecific pollen (HP) receipt (**A**) and conspecific pollen (CP) misplacement (**B**) and thus joint fitness gain. Paths reflect empirical tests of the hypotheses proposed in Fig. 1. The best supported model is presented (fig. S6). Black arrows indicate significant positive (solid) and negative (dashed) relationships. Arrow widths depict standardized coefficients. Attractive (red), female (brown) and male (purple) function traits are indicated, where ‘Dim’ indicates multivariate dimension from factor analysis of mixed data (figs. S2–S4).

To link functional traits to pollinator niche difference, we scored 20 floral traits (table S4) revealing substantial phenotypic variation among coexisting plants independent of abundance (Fig. 2B). We categorized these traits by function, that is, attractive (fig. S2), male (fig. S3) and female function (fig. S4), and performed multivariate analyses to obtain independent dimensions of trait variation within each functional group. We found multiple trait dimensions reflecting floral advertisement and mechanical fit to promote pollinator niche difference among coexisting plants. Specifically, the degree of pollination generalization, which was evolutionarily labile (Pagel’s λ = 0.075, *P* = 0.50; table S5), was significantly predicted by the first three dimensions of attractive traits (‘Dim’ 1–3, PSEM, *r* = 0.37, -0.32 and 0.22, respectively, *P* < 0.05; Fig. 3).

That is, species with open, funnelform (Dim 1) or aster-shaped (Dim 2) flowers were more generalized than pea (Dim 1) or salverform (Dim 2) flowers. Flower symmetry also predicted pollinator niche, with bilateral flowers more specialized than radial flowers (fig. S2), likely due to mechanical fit of pollinators (*20*). Similarly, longer flower tubes that filter pollinators by tongue length (*20*) were associated with less generalization. In addition, species with flowers horizontally arranged in inflorescences that increase advertisement size were more generalized than those with vertically arranged inflorescences or single flowers (fig. S2). In contrast to attractive traits, male function traits that potentially signal pollen rewards did not predict pollinator niche (Fig. 3).

To assess the fitness effects of pollinator niche difference and how it mediates rare species advantage, we taxonomically identified 3.1 million pollen grains that were deposited on stigmas of co-flowering species [*n* = 54 stigmas each of 66 species, see (*27*); table S6], over the two years of pollinator observations. We quantified male fitness loss as pollen misdelivered to coexisting heterospecifics (outgoing arrows, Fig. 2C), and female fitness loss due to HP receipt (incoming arrows) from the community-wide pollen transfer network, based on sufficient stigma sampling (fig. S5, rarefaction). Pollinator niche mediated fitness costs, with generalists receiving more HP and thus the potential for higher female fitness loss than specialists (*r* = 0.22, *P* < 0.05; Fig. 3A), in line with previous studies (*18, 23*). Although the positive relation of CP–HP receipt (*16, 18*) contributed to joint fitness gain (CP/ovule, *r* = 0.20, *P* < 0.05; Fig. 3A), pollinator diversity directly reduced joint fitness gain (*r* = -0.15, *P* < 0.05), possibly due to pollen consumption or other mechanisms of pollen loss during transport (*21*). As a result, there was a net negative fitness effect of generalization [*r* = -0.11 (-0.15 + 0.22 × 0.20); Fig. 3A], in line with the niche partitioning hypothesis (H1.1 net effect, Fig. 1C). Generalization, however, showed no direct effect on pollen misplacement (*r* = 0.01; Fig. 3B), in contrast to our hypothesis that it would lead to higher male fitness loss (H1.2, Fig. 1D). This suggests that pollinator diversity and visit quantity (*7*) are perhaps less important than other factors such as pollinator quality in mediating misplacement of pollen to heterospecifics. Taken together, these results demonstrated a fitness cost of generalization. In support of rare species advantage mediated by pollinator niche differentiation (H2 and H1.1, Fig. 1C), we found that rare species were more specialized (*r* = 0.41, *P* < 0.05) and thus had higher net joint fitness gain (Fig. 3A).

To test for asymmetric facilitation, we first differentiated abundance-based causes from trait-based causes of fitness differences among coexisting plants (Fig. 3). We found that different sets of traits influenced female and male fitness components, and these were distinct from the traits that influenced pollinator niche. Female function traits mediated fitness loss via HP receipt (Fig. 3A), while male function traits influenced fitness loss via CP misplacement (Fig. 3B). Larger stigmas and longer styles (*r* = 0.44, *P* < 0.05; Fig. 3A) or stigmas not extending beyond the corolla (*r* = -0.28, *P* < 0.05) led to higher HP receipt and thus female fitness loss. Pollen morphology explained a significant amount of variation among species in CP misplacement (*r* = 0.48 and -0.18 for Dim 3 and 2, respectively, *P* < 0.05; Fig. 3B). Small, textured (e.g. granulate or spiky) or regular-shaped (e.g. oval or round) pollen was more likely to be misplaced than pollen with opposite traits (i.e. large, smooth or irregular-shaped). In contrast to traits mediating fitness via pollen transport by pollinators (CP misplacement or HP receipt; Fig. 3), we detected little evidence for stigma–anther distance, a trait associated with self-pollen deposition [(*11*); Dim 4, fig. S4], to affect joint fitness gain (fig. S6). After accounting for the influence of floral functional traits, we isolated abundance-dependent effects on pollen transport and fitness and revealed evidence for facilitation. In line with the asymmetric facilitation hypothesis (Fig. 1C), rare species tended to receive more HP from coexisting heterospecifics than abundant species (albeit not statistically significantly more, *r* = -0.14; Fig. 3A). Yet pollinators that delivered HP also helped CP delivery (*16, 18*), contributing to a positive effect on joint fitness gain (*r* = 0.20, *P* < 0.05; Fig. 3A) which leads to a greater benefit for rare species. In addition, higher abundance led to greater male fitness loss via pollen misplacement (*r* = 0.62, *P* < 0.05; Fig. 3B), which translated into lowered joint fitness gain for abundant species (*r* = -0.19, *P* < 0.05), as hypothesized (H3.2, Fig. 1D). Overall, the results suggest that rare species experience mild increase in HP receipt but benefit from more HP due to the positive relation of CP–HP receipt, whereas abundant species suffer from greater pollen misplacement.

Our findings support the hypothesis that plant–pollinator interactions can favor rare species in species-rich, co-flowering communities, which contributes to the maintenance of plant diversity. Moreover, pollinator niche partitioning that leads to non-nestedness is essential for coexistence between rare and abundant species in this co-flowering community, where temporal niche partitioning is limited (*26*). When specialized plants do share some pollinator resources, asymmetric facilitation that benefits the rare at the cost of the abundant can serve as an alternative diversity-promoting mechanism. Our results underscore the potential to improve our understanding of the maintenance of rare species by considering not only seedling recruitment and growth but also community-wide interactions between plants and pollinators that affect fertilization success (*7, 8*). This is becoming more urgent than ever for predicting diversity maintenance as a result of anthropogenic changes in plant–pollinator mutualisms. In light of pollinator loss worldwide (*28*), overall diminished pollinator niche space may intensify plant competition and at the detriment of rare species that are more specialized than abundant species. Climatically induced shifts in plant abundance (*29*) may alter community-wide floral trait variation that is key to pollinator niche partitioning and subsequent pollen transport dynamics, and may also affect the strength of asymmetric facilitation if rare and abundant species respond differently to climate change. Understanding the mechanisms by which pollination contributes to the persistence of rare species is arguably one of the most critical tasks for biodiversity conservation in the Anthropocene.

## Acknowledgments

We thank J. Baker, D. Chang, A.M. Arters, U. Meenakshinathan, K. Doleski, S. Barratt-Boyes, R.A. Ashman and M. Holden for assistance with stigma pollen identification, floral trait measurements and insect specimen processing, J. Rawlins, J. Pawelek, R. Androw and B. Coulter for insect identification, J. Hyland and V. Verdecia for logistic support at Carnegie Museum of Natural History, and McLaughlin field station staff for logistic support of field work. We also thank the members of the Ashman, Wood and Turcotte laboratories for discussion.

## Author contributions

T.-L.A. conceived the study; N.W. and T.-L.A. led the conceptual development; N.W. analyzed the data and wrote the manuscript draft; T.-L.A., R.L.K. and G.A.-G. contributed to the final manuscript; R.L.K., E.M.O., R.A.H., G.A.-G. and T.-L.A. collected the data.

## Data and materials availability

All data are available in the main text or the supplementary materials.

## Supplementary Materials

### Materials and Methods

#### Study site and co-flowering community

The co-flowering community of the species-rich serpentine seep system at the McLaughlin Natural Reserve in California, USA (38.8582°N, 122.4093°W) is the subject of this study. The unique soil chemistry of these serpentine seeps and late summer moisture are important determinants of species that can survive in this environment, largely small herbaceous annuals and perennials (*30*). In this system, pollination is a strong force acting upon successful reproduction and likely coexistence, because seep drying restricts flowering and fruiting time, enforcing substantial flowering overlap and pollinator–meditated plant–plant interactions (*30–32*). We focused on a system that consisted of fives seeps (table S1), separated by 0.3–5 km (*31*). Each seep was visited once every week during the peak of flowering season (April–June) for a total of 9 to 10 weeks/year in 2016 and 2017.

#### Plant–pollinator interactions

Each week, plants at each seep were scored for plant–pollinator interactions. Observations were conducted between 0800–1700 h by two to three persons simultaneously. For the two species with crepuscular flowers (*Linanthus dichotomus* and *Chlorogalum pomeridianum*), pollinator observations were extended to 1900 h. All pollinators visiting a plant species were collected for identification with the exception of hummingbirds. We considered a legitimate plant–pollinator interaction only when a pollinator contacted the reproductive parts of a flower. Lepidopteran insects were preserved dry, and non-Lepidopteran insects were preserved with 100% ethanol in 1.5 mL microcentrifuge tubes in a -20 °C freezer until processed and pinned.

Pinned or ethanol preserved specimens were identified by experts: bees (Anthophila) by Jaime Pawelek (Wild Bee Garden Designs), beetles (Coleoptera) by Robert Androw (Carnegie Museum of Natural History, Pittsburgh, PA), flies (Diptera) by Ben Coulter (Carnegie Museum of Natural History, Pittsburgh, PA), and moths and butterflies (Lepidoptera), as well as remaining insects, by John Rawlins (Carnegie Museum of Natural History, Pittsburgh, PA). Insects were identified to the lowest taxonomical level possible (typically species level). All vouchered specimens were deposited at the Carnegie Museum of Natural History (Pittsburgh, PA).

We aimed to collect an equal number of pollinators per plant species (*n* = 150 on average) across seeps and years subject to plant availability. Rarefaction analysis (fig. S1) using the package iNEXT (*33*) in R v3.6.0 (*34*) showed that our sampling effort captured the majority of pollinator diversity for each of the 79 plant species (table S2).

#### Style collection

To characterize interspecific pollen transfer, styles from spent flowers of each species were collected on the same day as pollinator observations. Styles were collected from different individuals of each species in all cases except the very rare species. Three styles per species were stored together in a 1.5 mL microcentrifuge tube with 70% ethanol. For species with flowers that were too small to remove styles in the field (*n* = 9), we collected and stored whole flowers, and then styles were collected from these collected flowers in the lab with the aid of a dissecting microscope.

From this vast collection of styles, we employed a stratified random subsampling across all seeps and both years to achieve 90 (18 × 5) date–seep combinations, and a total of 54 styles per species for stigma pollen identification following the recommendation of (*35*). Fourteen species (*Achillea millefolium*, *Aquilegia eximia*, *Castilleja attenuata*, *Dichelostemma congestum*, *Eriogonum vimineum*, *Grindelia hirsutula*, *Hesperolinon californicum*, *Hemizonia congesta*, *Lomatium macrocarpum*, *Lupinus microcarpus*, *Micropus californicus*, *Microseris douglasii*, *Minuartia douglasii*, and *Vicia sativa*) did not meet our sampling goals and were excluded from downstream interspecific pollen transfer network and phylogenetic structural equation modeling (PSEM).

#### Floral abundance

Floral abundances were determined from weekly surveys of fixed plots (1 m × 3 m each) at each seep in both years (table S1). Plots were positioned along the length of each seep 1–20 m apart to capture plant species diversity within a site. At each sampling date, we recorded all open flowers for each species within a plot. For Asteraceae, we counted compact ‘heads’ as individual floral units. These surveys were carried out primarily during 1230–1400 h and were extended to 1900 h for species with crepuscular flowers. Floral abundance of each species was summed across fixed plots, seeps and years for downstream PSEM.

#### Floral functional traits

Flowers were collected from separate individuals of each species across one or more seeps, depending on rarity and stored in 70% ethanol. For 10 flowers per species, we measured 20 floral functional traits and subsequently categorized them according to their functions: attractive (*n* = 7), male (*n* = 8) and female (*n* = 5) function (figs. S2–S4).Three (*Adenostoma fasciculatum*, *Collomia diversifolia* and *Hoita macrostachya*) of the 72 species that were measured for floral functional traits (table S4) had no pollinator observations and were excluded from additional analyses.

The attractive traits included flower color, shape, symmetry, restrictiveness, inflorescence type, flower tube length and corolla limb length. Qualitative traits (flower color, shape, symmetry, restrictiveness, and inflorescence type) were scored in the field or from photographs of the species. Flower color was assessed based on human visual perception as white, yellow-orange, pink-red or purple. Flower shape was categorized as open, funnelform, labiate, salverform, pea- or aster-like, whereas symmetry was scored as radial or bilateral. We categorized restrictiveness (restrictive or not) based on whether morphological barriers that prevent some pollinators from accessing floral rewards exist or not, following (*36*). Inflorescence type was scored as single, horizontal or vertical cluster, where ‘single’ reflects flowers that are presented singly or widely spaced on a stem, and clusters are flowers arranged horizontally or vertically. Quantitative traits (flower tube and corolla limb length) were measured using preserved flower samples with a digital caliper (± 0.1 mm). Flower tube length was measured as the distance from the bottom of a superior ovary or the top of an inferior ovary to the top of corolla tube or petal separation. Tube length was scored as zero for species without flower tubes. Corolla limb length reflects the longest axis of corolla diameter. For Asteraceae, corolla limb length was defined as the average of a ray flower (tongue length) and disk flower (corolla diameter).

Male function traits included anther length, stamen number, height and exertion, and pollen shape, texture, area and width to length ratio. Anther length was measured along the longest axis. Stamen length was measured on the longest stamen of each species. Stamen number was counted directly (≤50) or estimated (if >50). Stamen exertion reflects the length that a stamen extends beyond (a positive value) or below (a negative value) the corolla and indicates stamen accessibility to pollinators. Pollen grains (*n* = 10 per species) were visualized under 400× magnification using a Leica DM500 microscope (Leica Microsystems, Wetzlar, Germany). Pollen shape was categorized as spherical, oval, or other (e.g. capsular, fenestrate, pyramidal or tetramerous shape). Pollen texture was categorized as psilate, granulate or spiky. Pollen area and width/length ratio were measured using ImageJ v1.47 (*37*) on photos of pollen grains. For Asteraceae, male function traits were obtained from hermaphroditic disk flowers only.

Female function traits included style length, stigma–anther separation, and stigma shape, area and exposure. Style length was the distance between ovary and stigma. Stigma–anther separation affects the potential for self-pollen deposition (*38*) and was measured as the distance between stigma and the closest anther. Stigma shape was categorized as lobed, non-lobed or ‘along’ the length of the style. We estimated stigma area from images of the stigmatic surface using ImageJ. Stigma exposure reflects the distance that a pistil extends beyond corolla (a positive value) or below corolla (a negative value). For Asteraceae, female function traits were averaged between female ray flowers and hermaphroditic disk flowers.

#### Pollen identification on stigmas

To taxonomically identify pollen grains on stigmas, we created a pollen library for all flowering plant species in the seeps (R. Hayes, N. Cullen, R. Kaczorowski, and T-L. Ashman, in prep). Species-specific pollen traits (i.e. size, shape, texture and aperture numbers) were obtained for acetolyzed pollen (*39*) collected from anthers of flowers in the seeps. Flowers from non-focal plants outside the seeps (e.g. grasses and trees) were also collected to facilitate the identification of pollen communities received by individual stigmas.

To characterize conspecific (CP) and heterospecific pollen (HP) received by stigmas, we acetolyzed on average 54 styles (range = 36–57) from each species sampled in a stratified random manner across all seeps and years, as described above (in section *Style collection*). Specifically, we acetolyzed the contents (*39*) of each sample tube (3 styles and their pollen grains) to achieve a volume of 20 µL. We then enumerated pollen of a 5 µL aliquot using a hemocytometer and calculated the total amount of pollen grains per style. When pollen was too dense to count, we diluted the 20 µL to 100 or 200 µL and adjusted final counts accordingly.

Each pollen grain was identified to species (including both CP and HP) based on the pollen library. When pollen was from species not present in the pollen library, we designated it as from a specific unknown species (e.g. U1, U2, etc.). But in the cases where we were unable to distinguish a pollen grain among congeners or morphologically similar species, we assigned it to a congener- or morphospecies-group. We then used a fractional identity approach to assign pollen grains within these groups. Fractional identity was based on relative probabilities as a function of floral abundance at the sampling seep and date. To examine how fractional identity influenced the estimate of HP on stigmas, we compared HP richness when fractional identities were excluded and included. A strong positive correlation between the two approaches was observed (*r* = 0.73; fig. S5A), supporting the use of fractional identity. Rarefaction analysis of pollen grains (with fractional identity) showed that our sampling effort of styles captured the majority of HP donor species for each recipient species (fig. S5 B and C).

To account for variation in the number of sampled styles among species, we standardized pollen data to the same number of stigmas (*n* = 54) across species, which outperforms standardization based on rarefaction (*40*) (i.e. the same minimum number of stigmas across species). The standardized pollen data were used for subsequent analyses in constructing the interspecific pollen transfer network.

#### Plant phylogeny

The phylogenetic tree of all 79 co-flowering species was constructed based on PhytoPhylo megaphylogeny of vascular plants (*41, 42*) and the Open Tree of Life (*43*) using the R packages ape (*44*), rotl (*45*) and phytools (*46*). Specifically, we used genus-level megaphylogeny as the backbone. For *Githopsis* and *Hemizonia* that were not present in the megaphylogeny, we determined their positions in the tree according to the Open Tree of Life and branch lengths according to family-level megaphylogeny. For all congeners, we obtained their relative positions and branch lengths from megaphylogeny, the Open Tree of Life or published genus-specific trees (*47–53*). In the cases where branch length was not available for a species (*Astragalus rattanii*, *Calochortus amabilis*, *Castilleja attenuate*, *Castilleja rubicundula*, *Mimulus layneae*, and *Trifolium obtusiflorum*), we used the phylogenetic information of the closest relative within the same genus as a surrogate from the aforementioned sources. The phylogenetic tree was visualized using the package ape.

#### Multivariate analyses of floral traits

To examine floral trait variation among co-flowering species, we used trait mean averaged across the 10 flowers each species. We first assessed the overall floral trait variation by performing a factor analysis of mixed data (FAMD) of all the 20 quantitative and qualitative traits using the package FactoMineR (*54*), as shown in Fig. 2B. Instead of imputation, missing data were omitted from the FAMD. We then performed FAMD for attractive, male and female function traits independently, and used the first three dimensions from each for subsequent PSEM. Our choice of the first three dimensions aimed to capture a large amount of trait variation (48%–71% here) while avoiding overparameterizing PSEM. It is worth noting that different from multivariate analyses of quantitative data (e.g. principle component analysis), in FAMD the same qualitative trait can contribute to more than one independent FAMD dimension, because of multiple (>2, not binary) categories within each qualitative trait.

#### Plant–pollinator network

We constructed the network based on 7324 total plant–pollinator interactions observed across all seeps in both years using the package bipartite (*55*). To evaluate pollinator niche partitioning, we assessed whether co-flowering plant species were limited in pollinator sharing, that is, more specialized than expected by random interactions with pollinator partners, at the species, group (of plants as a whole) and network levels using multiple metrics. At the species level, we used the metrics of 1) pollinator Shannon diversity, which considers both pollinator richness and interaction frequencies, and 2) similarity between pollinator use and availability [i.e. proportional similarity, (*56*)], which indicates increased generalization when it increases from 0 to 1. At the group level, we used the metrics of 3) mean number of shared pollinator partners between any two plants, and 4) mean similarity in pollinator assemblage between any two plants [i.e. niche overlap using Horn’s index in bipartite; (*57*)], which considers both pollinator identity and interaction frequencies. At the network level, we used 5) NODF [nestedness overlap and decreasing fill, (*58*)] and 6) its weighted version that takes into account interaction frequencies. Both unweighted and weighted NODF reflect specialization asymmetry (i.e. specialists interacting with the subset of pollinator partners of generalists), with increased nestedness when the metrics increase from 0 to 100. For the next step, we compared these observed metrics to null models. We constructed the null model by rewiring interactions while keeping total interaction frequencies of individual plants and pollinators constant [i.e. r2dtable algorithm; (*59*)] using the packages bipartite and vegan (*60*). Based on 1000 random replications of the null model, we obtained null distributions of individual metrics, and calculated null mean and the 95% confidence intervals (i.e. the 2.5th and 97.5th percentiles). Statistical significance of each observed metric was obtained by comparing the observed value to the null confidence intervals. The two-sided *P* value was calculated as how often the observed metric was greater or smaller than all 1000 random replicates.

The plant–pollinator network was visualized as an interaction matrix using the package ggplot2 (*61*). Plant species were arranged according to their phylogenetic positions. Pollinator species were arranged according to the number and community assemblage of the plant species that they visited, the latter of which was obtained using the first axis of a canonical correspondence analysis (CCA) in the package vegan.

#### Interspecific pollen transfer network and pollination-mediated fitness

A pollen transfer network that describes pollinator-mediated pollen delivery among co-flowering plants was constructed using the package igraph (*62*), based on standardized pollen receipt by the same number of stigmas (*n* = 54) across species using fractional identity as described above (see section *Pollen identification on stigmas*). The standardized pollen receipt can remove variation in sampling effort and effectively reflect per capita estimates of pollen received and donated for each species. In this directed pollen transfer network, arrows link pollen donor to recipient species. For a focal species, the number of incoming arrows represent the number of heterospecific species from which the focal species receives pollen. Arrow widths represent the amount of pollen from individual donors, the sum of which (i.e. strength-in or total HP receipt) indicates female fitness loss for that species. In contrast, outgoing arrows indicate the number of heterospecific species to which a focal species donates pollen, and the amount of conspecific pollen that is misdelivered to coexisting heterospecifics (i.e. strength-out) indicates male fitness loss. Pollen transfer network was visualized using Gephi v0.9.2 (*63*).

The successful pollination outcome of pollen transport is CP deposition on conspecific stigmas. This is a per capita estimate of joint male and female fitness as it represents their mating success (via male and female gametes), based on the standardized pollen receipt data. We normalized this estimate of joint fitness gain by accounting for the differences across species in female gametes (i.e. ovule number), that is, CP per ovule. Ovule number was obtained from field-collected, ethanol-preserved flowers that were collected for floral trait measurements (see section *Floral functional traits*) or from our previous greenhouse studies (*Mimulus layneae*, *Mimulus guttatus* and *Mimulus nudatus*) (*64*).

#### Phylogenetic signal

To account for evolutionary dependence among co-flowering species in SEM, we first examined the phylogenetic signals of floral functional traits [20 traits as well as the first three FAMD dimensions for each of the functional groups of traits (attractive, male and female)], pollinator niche breath (species-level metrics from the plant–pollinator network) and pollination-mediated fitness estimates (HP receipt, CP misplacement and CP/ovule). For continuous variables, we assessed the phylogenetic signals using Pagel’s λ (*65*) implemented in the package phylosignal (*66*). Pagel’s λ informs whether variables of interest have evolved independently of phylogeny (λ = 0, a lack of phylogenetic signal), or under a Brownian motion process (i.e. random drift; λ = 1) or other processes (0 < λ < 1). For categorical variables, we converted variables with more than two categories into binary variables using dummy coding (i.e. 1 for a focal category and 0 for non-focal categories), and examined phylogenetic signals using the *D* statistic (*67*) in the package caper (*68*). In contrast to Pagel’s λ, *D* = 1 indicates the lack of phylogenetic signal, and *D* = 0 indicates Brownian motion evolution. *D* > 1 or *D* < 0 signifies phylogenetic overdispersion and clustering, respectively. In contrast to pollinator niche breath and pollination-mediated fitness (table S5), floral functional traits including FAMD dimensions exhibited strong phylogenetic signals (table S5). Such evolutionary dependence was considered in subsequent PSEM.

#### Phylogenetic structural equation modeling (PSEM)

We conducted PSEM to create a process-based hypothesis for the links between floral functional traits and plant rarity (abundance) to pollination niche breath (generalization) and pollination-mediated fitness losses and gains, and thereby could test explicitly the paths associated with niche partitioning or asymmetric facilitation (Fig. 1). We used the package phylopath (*69*) with Pagel’s model to account for evolutionary dependence among species. For pollination generalization at the species level, we used pollinator Shannon diversity as described above, because this metric considered both pollinator richness and interaction frequencies and showed a strong correlation with the other species-level metric here (i.e. similarity between pollinator use and availability, *r* = 0.79, *P* < 0.001). We checked for collinearity among predictors, and confirmed low correlations among trait dimensions from the three functional categories (all *r* < 0.4), using the package psych (*70*). To improve normality in linear models of PSEM, we power transformed endogenous variables if necessary, with the optimal power parameter determined using the Box–Cox method in the package car (*71*). Specifically, natural logarithm transformation was applied to joint fitness gain (CP/ovule) and female fitness loss (HP receipt or strength-in), and the optimal power parameter was 0.2 for male fitness loss (CP misplacement or strength-out).

To account for the potential influence of self pollen deposition in contrast to pollinator-mediated pollen deposition, we built two models (fig. S6) that considered pollinator-mediated pollen deposition alone (model 1) and both pollinator-mediated and self pollen deposition (model 2) for evaluating niche partitioning and asymmetric facilitation. In model 1, joint fitness gain (CP/ovule) was hypothesized to be influenced by pollinator niche (generalization) and pollinator-mediated HP receipt (Fig. 1C and fig. S6A) and CP misplacement (Fig. 1D and fig. S6B). Relative to model 1, model 2 added the possibility that stigma–anther distance (i.e. herkogamy) may influence self pollen deposition and thus CP/ovule. We compared the two models using the *C* statistic Information Criterion [CIC, (*72*)], which penalizes model fit by parameter numbers in PSEM (*72, 73*). Following (*72*), the best-supported model produces the lowest CIC and its difference with other models (ΔCIC) is greater than 2.

The standard errors and 95% confidence intervals of individual standardized regression coefficients (of links, *r*) were obtained via bootstrapping (*n* = 1000). When the 95% confidence intervals did not overlap with zero, a link was considered significant. Using significant links, we assessed whether the proposed hypotheses (Fig. 1 C and D) were supported. Specifically, to support the hypothesis of the fitness costs of generalization (H1.1 and H1.2 net effects), we required generalization being directly negatively linked to joint fitness gain, and/or indirectly negatively linked to joint fitness gain via influencing female fitness loss (Fig. 1C, H1.1) and male fitness loss (Fig. 1D, H1.2). To support the hypothesis of rare species advantage due to pollinator niche partitioning, we required that abundance was positively linked to generalization (H2) and generalization was negatively linked to joint fitness gain (i.e. fitness costs of generalization; H1.1 and H1.2 net effects), leading to higher joint fitness gain of rare species than abundant species. To support the hypothesis of asymmetric facilitation, first we required that rare species were facilitated more than abundant species by receiving more CP along with accompanying HP (i.e. abundance negatively linked to HP receipt), leading to increased joint fitness gain (i.e. HP receipt positively linked to CP/ovule; Fig. 1C, H3.1). Second, we required that abundant species as facilitators experienced higher male fitness loss (i.e. abundance positively linked to CP misplacement), leading to reduced joint fitness gain relative to rare species (i.e. CP misplacement negatively linked to CP/ovule; Fig. 1D, H3.2).

---

**Fig. S1.**
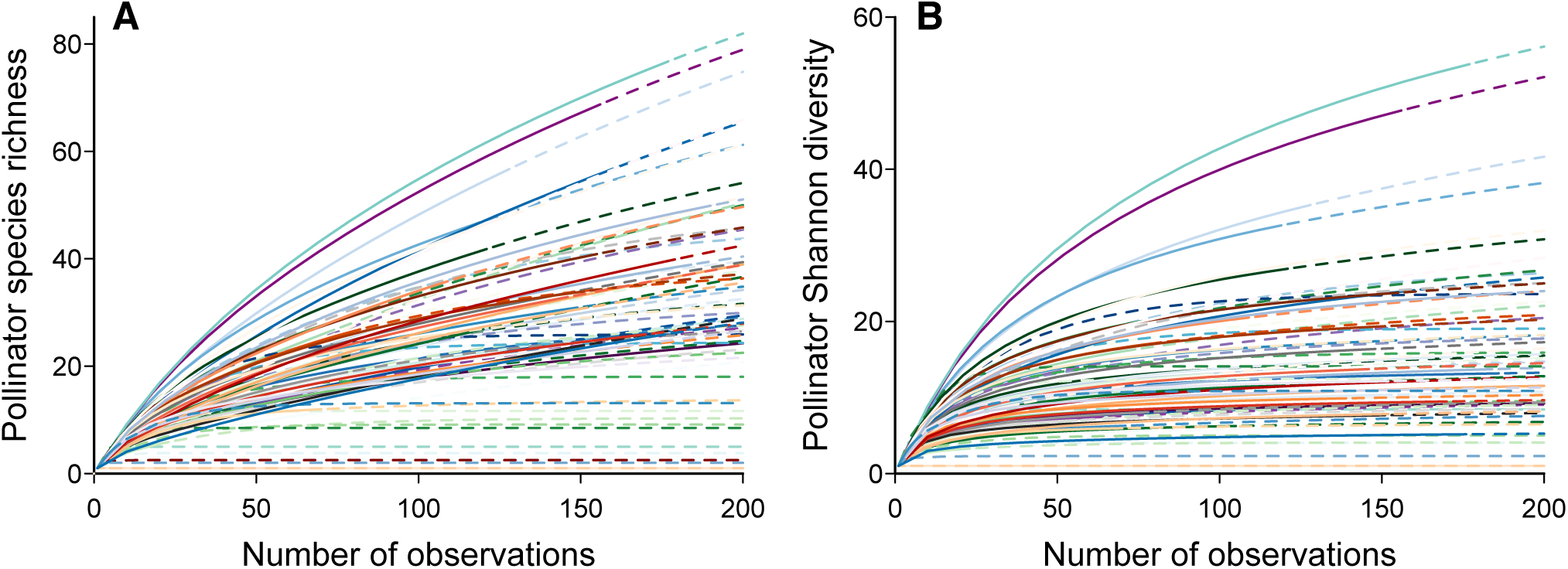
Rarefaction shows that the majority of pollinator diversity was captured with our sampling intensity. The number of pollinators (*x*-axis) observed for each plant species is represented by the solid portion of each colored line, whereas the dashed portion indicates extrapolation in the rarefaction analysis using the R package iNEXT (*33*). Both pollinator species richness (**A**) and Chao’s Shannon diversity (**B**) started to level off at the observed number of pollinators for most plant species, reflecting sufficient sampling to capture pollinator diversity.

**Fig. S2.**
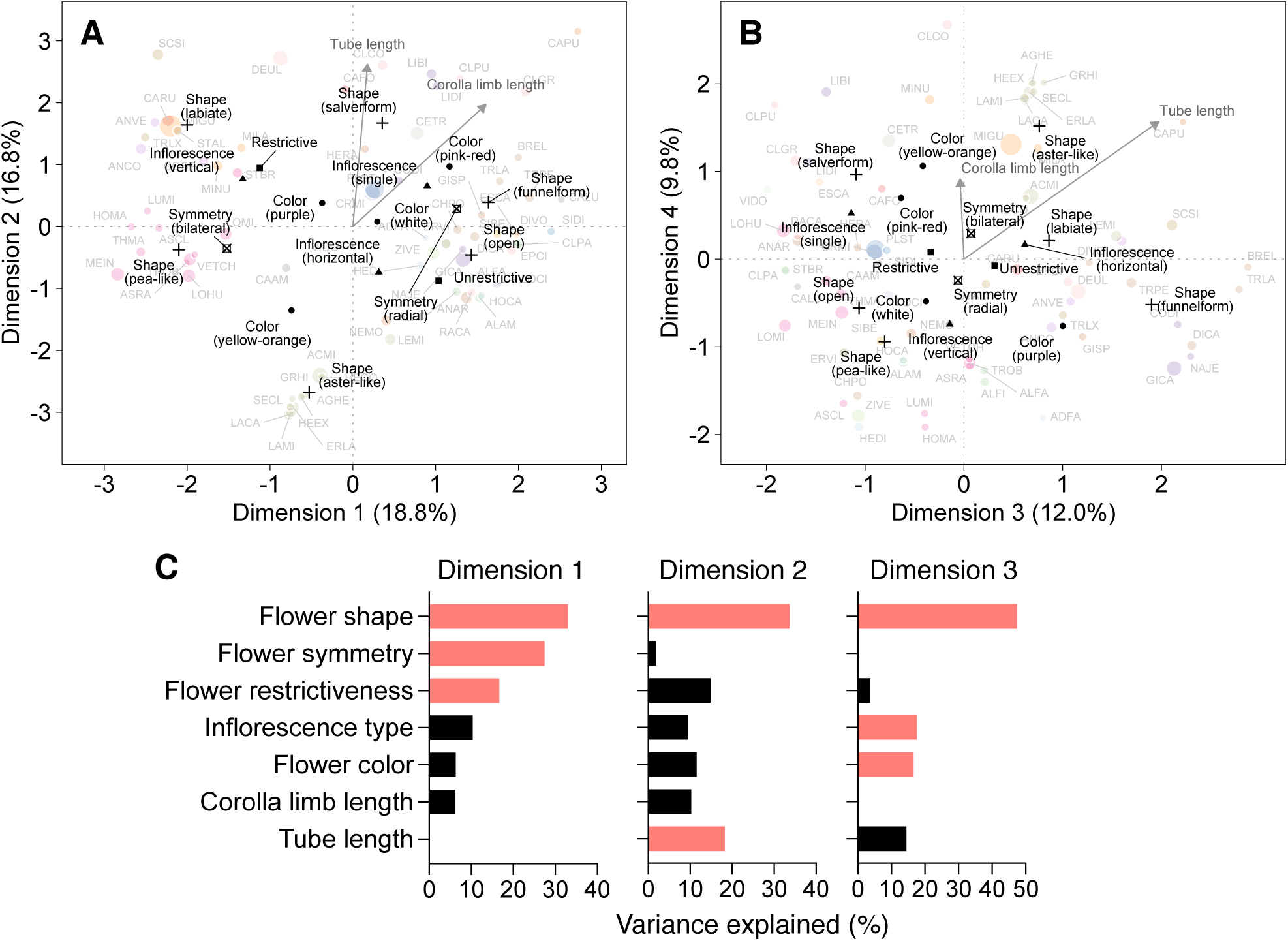
Multivariate analysis of floral traits associated with pollinator attraction. In the first four dimensions (**A** and **B**) of the factor analysis of mixed data (FAMD), the centroid of each category within a qualitative trait is indicated, with symbol shape representing different qualitative traits. Quantitative traits are represented by arrows. Individual plant species are shown in the background with colors indicating plant family and symbol sizes indicating relative abundances. (**C**) The traits that contributed to ≥15% of variation of the first three dimensions are highlighted in color.

**Fig. S3.**
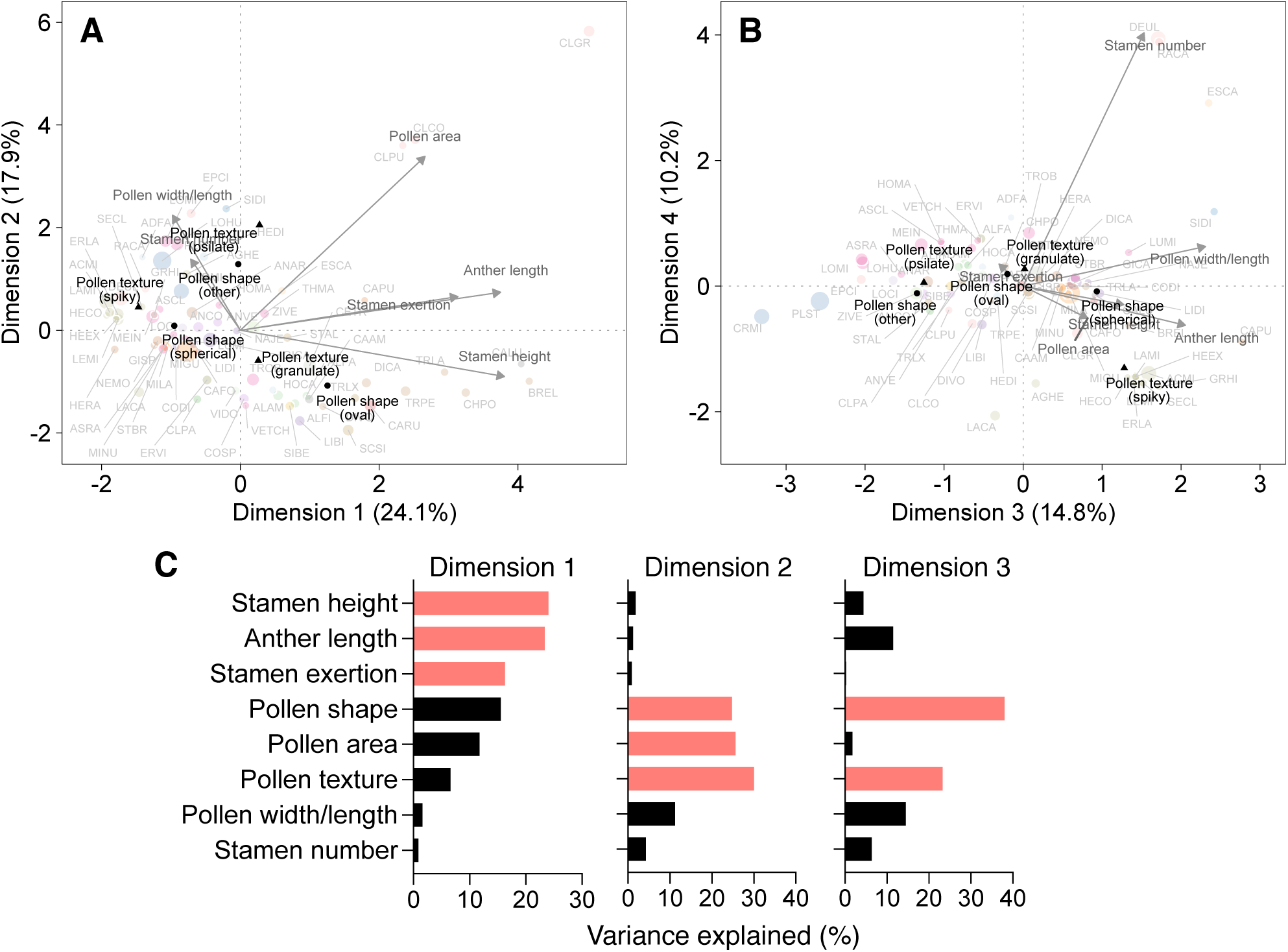
Multivariate analysis of traits associated with male function. In the first four dimensions (**A** and **B**) of the factor analysis of mixed data (FAMD), the centroid of each category within a qualitative trait is indicated, with symbol shape representing different qualitative traits. Quantitative traits are represented by arrows. Individual plant species are shown in the background with colors indicating plant family and symbol sizes indicating relative abundances. (**C**) The traits that contributed to ≥15% of variation of the first three dimensions are highlighted in color.

**Fig. S4.**
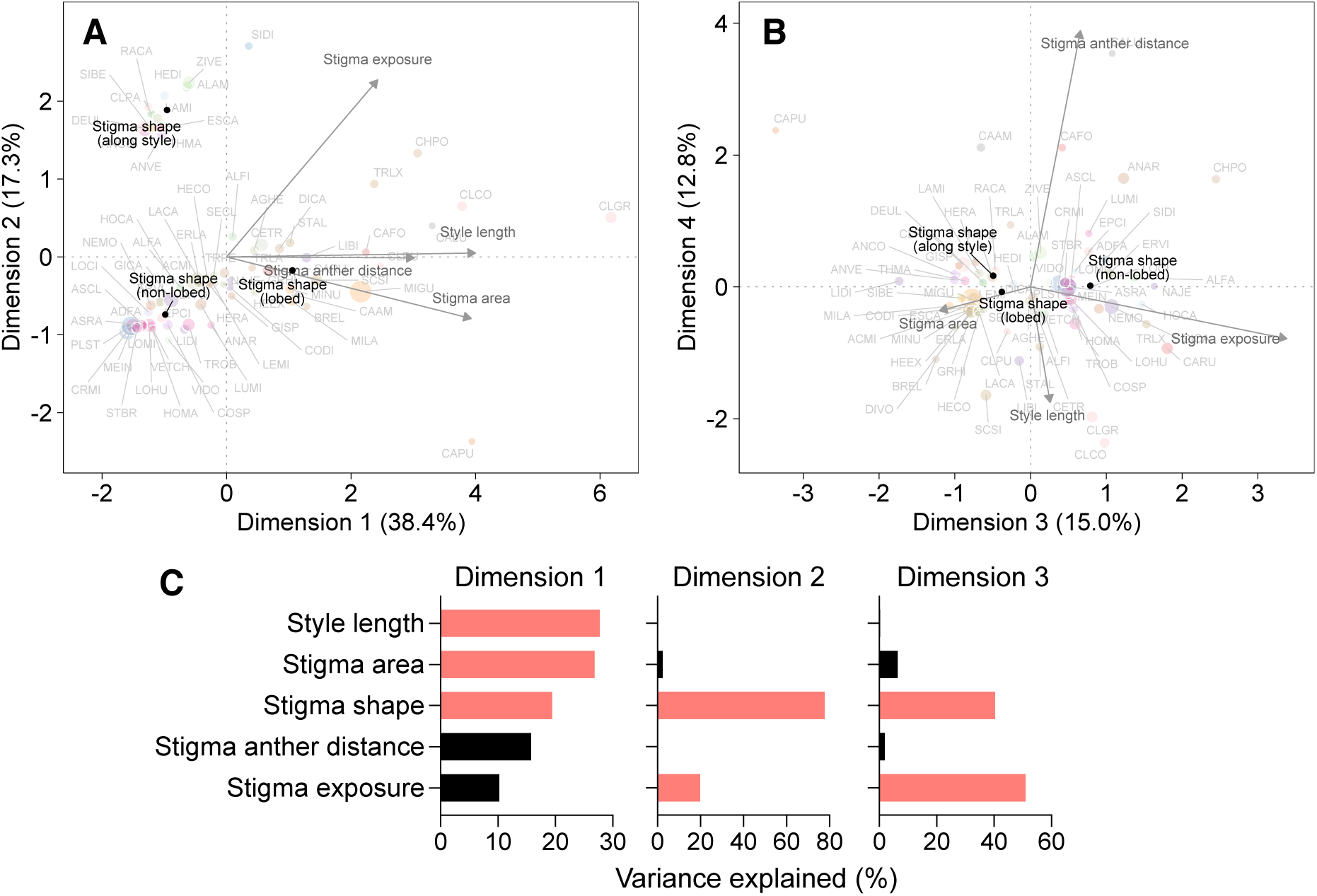
Multivariate analysis of traits associated with female function. In the first four dimensions (**A** and **B**) of the factor analysis of mixed data (FAMD), the centroid of each category within a qualitative trait is indicated, with symbol shape representing different qualitative traits. Quantitative traits are represented by arrows. Individual plant species are shown in the background with colors indicating plant family and symbol sizes indicating relative abundances. (**C**) The traits that contributed to ≥15% of variation of the first three dimensions are highlighted in color.

**Fig. S5.**
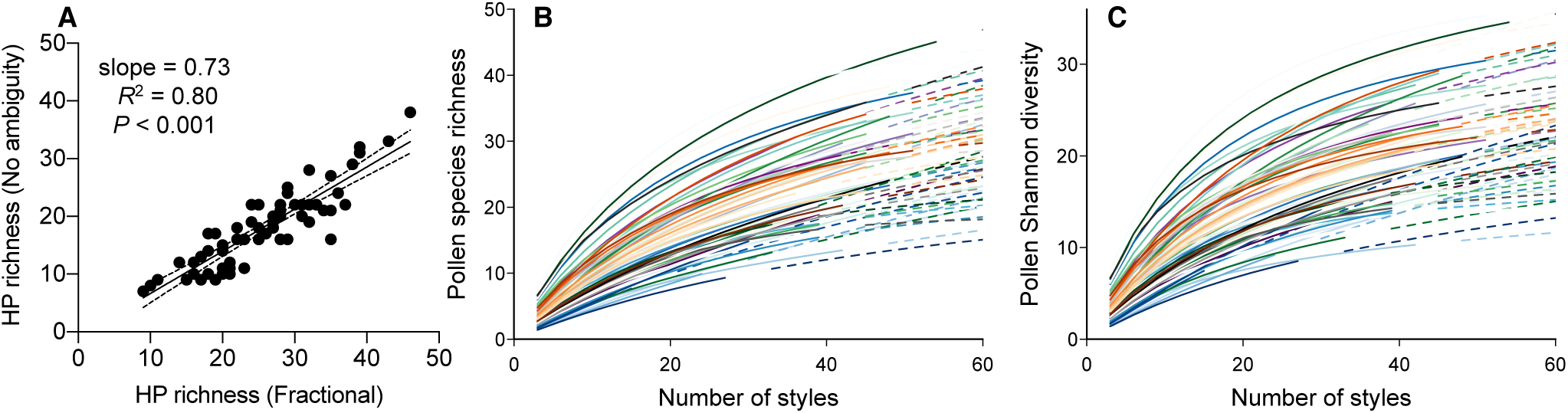
Validation of fractional identity approach and rarefaction of pollen received by stigmas. (**A**) There was a strong correlation (with dotted 95% confidence intervals) of heterospecific pollen (HP) richness when fractionally identified pollen grains were excluded (*y*-axis, ‘no ambiguity’) and included (*x*-axis, ‘fractional’). Rarefaction analysis using the R package iNEXT (*33*) showed that the majority of pollen species richness (**B**) and Chao’s Shannon diversity (**C**) were captured by the sampled styles (*n* = 54 on average) for each plant species (individual colored lines). The observed (solid) and extrapolated (dashed) portion of each rarefaction line are indicated.

**Fig. S6.**
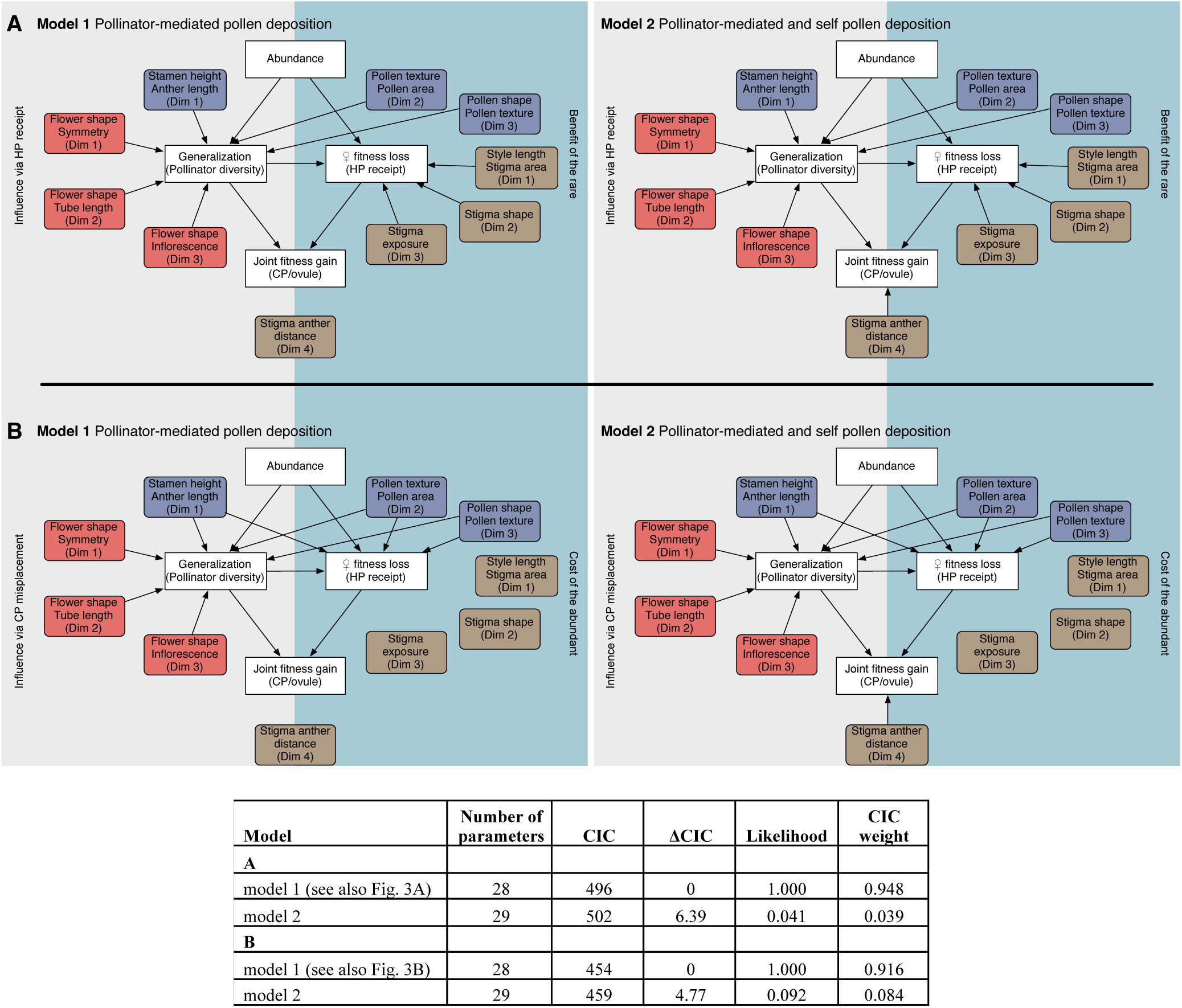
Phylogenetic structural equation models. Two models that considered pollinator-mediated pollen deposition alone (model 1) and both pollinator-mediated and self pollen deposition (model 2) were included for evaluating niche partitioning and asymmetric facilitation via HP receipt (**A**) and CP misplacement (**B**). Relative to model 1, model 2 added the possibility that stigma–anther distance may influence self pollen deposition and thus CP/ovule. Model 1 in both A and B represent the best-supported models and are presented in Fig. 3. CIC, the *C* statistic Information Criterion.

**Table S1.** Serpentine seep system within the McLaughlin Nature Reserve study area. Pollinators, flower abundances and styles were collected across this system. Within each seep, flower abundances were recorded 9*–10 times each year across 8–16 1 m × 3 m plots per seep, depending on seep size and flowering duration.

| **Seep** | **Seep name** | **Latitude** | **Longitude** | **Seep area (m<sup>2</sup>)** | **Number of survey plots in 2016** | **Number of survey plots in 2017** |
| --- | --- | --- | --- | --- | --- | --- |
| BS | Banana Slug | 38.8622°N | 122.3992°W | 2200 | 13* | 13 |
| RHA | Research Hill A | 38.8589°N | 122.4103°W | 2200 | 12 | 11 |
| RHB | Research Hill B | 38.8572°N | 122.4075°W | 2200 | 16 | 15 |
| TPW | Tailings Pond West | 38.8661 N | 122.4511°W | 2300 | 11 | 11 |
| TP9 | Tailings Pond Site 9 | 38.8639°N | 122.4272°W | 1300 | 9* | 8 |

**Table S3.** Plant–pollinator network metrics and comparisons to null models. Null mean, 95% confidence intervals (CLs), and *P* values were obtained from 1000 random replications of the ‘r2dtable’ null model. Plant species (abbreviated as the first two letters of genus and species names) are indicated. NODF, nestedness overlap and decreasing fill.

| **Metric** | **Observed value** | **Null mean** | **CL lower** | **CL upper** | ***P*** |
| --- | --- | --- | --- | --- | --- |
| **Network level** |  |  |  |  |  |
| NODF | 13.00 | 28.78 | 28.13 | 29.45 | 0.00 |
| Weighted NODF | 5.86 | 12.38 | 11.97 | 12.78 | 0.00 |
| **Group level** |  |  |  |  |  |
| Mean shared partners | 3.31 | 17.28 | 16.88 | 17.71 | 0.00 |
| Niche overlap Horn’s index | 0.09 | 0.41 | 0.40 | 0.43 | 0.00 |
| **Species level** |  |  |  |  |  |
| Pollinator Shannon diversity |  |  |  |  |  |
| ACMI | 0.00 | 0.00 | 0.00 | 0.00 | 0.00 |
| AGHE | 2.34 | 4.05 | 3.88 | 4.21 | 0.00 |
| ALAM | 3.57 | 3.97 | 3.77 | 4.15 | 0.00 |
| ALFA | 2.71 | 3.26 | 3.00 | 3.46 | 0.00 |
| ALFI | 3.49 | 3.93 | 3.75 | 4.11 | 0.00 |
| ANAR | 1.84 | 3.67 | 3.44 | 3.86 | 0.00 |
| ANCO | 2.58 | 4.13 | 3.97 | 4.29 | 0.00 |
| ANVE | 2.16 | 4.01 | 3.82 | 4.18 | 0.00 |
| AQEX | 0.00 | 1.07 | 0.64 | 1.10 | 0.00 |
| ASCL | 1.73 | 3.70 | 3.46 | 3.89 | 0.00 |
| ASRA | 0.95 | 1.56 | 1.33 | 1.61 | 0.00 |
| BREL | 2.44 | 3.87 | 3.65 | 4.05 | 0.00 |
| CAAM | 1.10 | 1.07 | 0.64 | 1.10 | 0.13 |
| CAAT | 0.69 | 0.68 | 0.69 | 0.69 | 0.05 |
| CAFO | 0.00 | 1.35 | 1.04 | 1.39 | 0.00 |
| CALU | 3.00 | 3.71 | 3.49 | 3.90 | 0.00 |
| CAPU | 1.83 | 3.88 | 3.67 | 4.07 | 0.00 |
| CARU | 2.61 | 3.82 | 3.61 | 4.01 | 0.00 |
| CETR | 2.11 | 4.12 | 3.95 | 4.28 | 0.00 |
| CHPO | 2.72 | 4.08 | 3.90 | 4.25 | 0.00 |
| CLCO | 2.66 | 4.07 | 3.90 | 4.23 | 0.00 |
| CLGR | 2.62 | 4.12 | 3.95 | 4.29 | 0.00 |
| CLPA | 2.55 | 3.38 | 3.14 | 3.59 | 0.00 |
| CLPU | 2.61 | 3.47 | 3.25 | 3.66 | 0.00 |
| COSP | 2.01 | 3.61 | 3.39 | 3.81 | 0.00 |
| CRMI | 3.87 | 4.05 | 3.88 | 4.22 | 0.04 |
| DEUL | 2.24 | 4.21 | 4.06 | 4.36 | 0.00 |
| DICA | 2.89 | 3.73 | 3.50 | 3.93 | 0.00 |
| DICO | 1.56 | 1.74 | 1.56 | 1.79 | 0.03 |
| DIVO | 1.75 | 3.07 | 2.79 | 3.27 | 0.00 |
| EPCI | 2.80 | 3.69 | 3.45 | 3.89 | 0.00 |
| ERLA | 3.99 | 4.11 | 3.95 | 4.26 | 0.15 |
| ERVI | 1.97 | 2.27 | 1.97 | 2.40 | 0.03 |
| ESCA | 2.86 | 4.04 | 3.86 | 4.20 | 0.00 |
| GICA | 3.18 | 4.07 | 3.88 | 4.23 | 0.00 |
| GISP | 2.30 | 2.55 | 2.27 | 2.71 | 0.05 |
| GITR | 0.69 | 0.68 | 0.69 | 0.69 | 0.03 |
| GRHI | 2.57 | 3.84 | 3.64 | 4.04 | 0.00 |
| HECA | 1.20 | 2.93 | 2.65 | 3.12 | 0.00 |
| HECO | 1.23 | 2.36 | 2.09 | 2.48 | 0.00 |
| HEDI | 2.17 | 3.85 | 3.64 | 4.04 | 0.00 |
| HEEX | 2.48 | 3.18 | 2.93 | 3.39 | 0.00 |
| HERA | 1.39 | 1.35 | 1.04 | 1.39 | 0.20 |
| HOCA | 2.46 | 3.94 | 3.75 | 4.12 | 0.00 |
| LACA | 3.30 | 3.94 | 3.75 | 4.12 | 0.00 |
| LAMI | 2.68 | 4.10 | 3.91 | 4.25 | 0.00 |
| LEMI | 2.31 | 4.06 | 3.89 | 4.22 | 0.00 |
| LIBI | 1.95 | 3.53 | 3.30 | 3.73 | 0.00 |
| LIDI | 2.37 | 2.86 | 2.58 | 3.09 | 0.00 |
| LOCI | 2.09 | 3.73 | 3.50 | 3.93 | 0.00 |
| LOHU | 2.80 | 4.04 | 3.85 | 4.22 | 0.00 |
| LOMA | 0.00 | 0.68 | 0.69 | 0.69 | 0.00 |
| LOMI | 2.04 | 4.04 | 3.85 | 4.21 | 0.00 |
| LUMI | 0.00 | 0.00 | 0.00 | 0.00 | 0.00 |
| MEIN | 2.22 | 3.28 | 3.00 | 3.49 | 0.00 |
| MICA | 0.00 | 0.00 | 0.00 | 0.00 | 0.00 |
| MICDO | 0.00 | 1.35 | 1.04 | 1.39 | 0.00 |
| MIGU | 2.50 | 4.28 | 4.14 | 4.41 | 0.00 |
| MILA | 2.29 | 4.01 | 3.82 | 4.18 | 0.00 |
| MINDO | 0.69 | 0.68 | 0.69 | 0.69 | 0.03 |
| MINU | 2.85 | 3.75 | 3.54 | 3.95 | 0.00 |
| NAJE | 2.96 | 4.04 | 3.86 | 4.21 | 0.00 |
| NEMO | 3.17 | 4.05 | 3.89 | 4.22 | 0.00 |
| PLST | 3.33 | 3.97 | 3.77 | 4.15 | 0.00 |
| RACA | 2.81 | 3.88 | 3.68 | 4.07 | 0.00 |
| SCSI | 1.79 | 3.67 | 3.43 | 3.86 | 0.00 |
| SECL | 2.06 | 4.05 | 3.87 | 4.22 | 0.00 |
| SIBE | 3.02 | 3.90 | 3.69 | 4.08 | 0.00 |
| SIDI | 2.62 | 4.03 | 3.86 | 4.20 | 0.00 |
| STAL | 2.25 | 4.09 | 3.91 | 4.26 | 0.00 |
| STBR | 2.52 | 4.11 | 3.95 | 4.27 | 0.00 |
| THMA | 0.69 | 0.68 | 0.69 | 0.69 | 0.04 |
| TRLA | 3.10 | 3.85 | 3.63 | 4.05 | 0.00 |
| TRLX | 2.50 | 4.05 | 3.87 | 4.21 | 0.00 |
| TROB | 2.41 | 4.08 | 3.90 | 4.26 | 0.00 |
| TRPE | 3.17 | 4.13 | 3.97 | 4.29 | 0.00 |
| VETCH | 0.69 | 1.34 | 1.04 | 1.39 | 0.00 |
| VIDO | 1.68 | 2.10 | 1.83 | 2.20 | 0.01 |
| ZIVE | 1.64 | 4.11 | 3.94 | 4.28 | 0.00 |
| Proportional similarity |  |  |  |  |  |
| ACMI | 0.01 | 0.02 | 0.00 | 0.07 | 0.99 |
| AGHE | 0.20 | 0.61 | 0.56 | 0.65 | 0.00 |
| ALAM | 0.20 | 0.58 | 0.53 | 0.62 | 0.00 |
| ALFA | 0.18 | 0.37 | 0.29 | 0.43 | 0.00 |
| ALFI | 0.33 | 0.57 | 0.52 | 0.61 | 0.00 |
| ANAR | 0.17 | 0.48 | 0.42 | 0.54 | 0.00 |
| ANCO | 0.26 | 0.64 | 0.59 | 0.68 | 0.00 |
| ANVE | 0.21 | 0.59 | 0.55 | 0.64 | 0.00 |
| AQEX | 0.02 | 0.06 | 0.01 | 0.13 | 0.23 |
| ASCL | 0.12 | 0.49 | 0.43 | 0.55 | 0.00 |
| ASRA | 0.01 | 0.09 | 0.02 | 0.17 | 0.00 |
| BREL | 0.26 | 0.54 | 0.49 | 0.59 | 0.00 |
| CAAM | 0.00 | 0.06 | 0.01 | 0.12 | 0.00 |
| CAAT | 0.07 | 0.04 | 0.00 | 0.10 | 0.31 |
| CAFO | 0.02 | 0.07 | 0.01 | 0.14 | 0.11 |
| CALU | 0.24 | 0.49 | 0.44 | 0.55 | 0.00 |
| CAPU | 0.21 | 0.55 | 0.50 | 0.60 | 0.00 |
| CARU | 0.24 | 0.53 | 0.48 | 0.58 | 0.00 |
| CETR | 0.32 | 0.63 | 0.60 | 0.67 | 0.00 |
| CHPO | 0.32 | 0.62 | 0.58 | 0.66 | 0.00 |
| CLCO | 0.28 | 0.61 | 0.57 | 0.66 | 0.00 |
| CLGR | 0.29 | 0.63 | 0.59 | 0.67 | 0.00 |
| CLPA | 0.11 | 0.40 | 0.33 | 0.47 | 0.00 |
| CLPU | 0.13 | 0.43 | 0.36 | 0.49 | 0.00 |
| COSP | 0.17 | 0.47 | 0.40 | 0.52 | 0.00 |
| CRMI | 0.21 | 0.61 | 0.57 | 0.65 | 0.00 |
| DEUL | 0.17 | 0.67 | 0.63 | 0.70 | 0.00 |
| DICA | 0.24 | 0.50 | 0.45 | 0.55 | 0.00 |
| DICO | 0.03 | 0.10 | 0.03 | 0.18 | 0.05 |
| DIVO | 0.17 | 0.32 | 0.24 | 0.39 | 0.00 |
| EPCI | 0.20 | 0.49 | 0.43 | 0.55 | 0.00 |
| ERLA | 0.35 | 0.63 | 0.59 | 0.67 | 0.00 |
| ERVI | 0.06 | 0.17 | 0.09 | 0.26 | 0.01 |
| ESCA | 0.30 | 0.60 | 0.56 | 0.64 | 0.00 |
| GICA | 0.36 | 0.61 | 0.57 | 0.66 | 0.00 |
| GISP | 0.12 | 0.22 | 0.13 | 0.30 | 0.02 |
| GITR | 0.02 | 0.04 | 0.00 | 0.11 | 0.53 |
| GRHI | 0.16 | 0.54 | 0.49 | 0.59 | 0.00 |
| HECA | 0.17 | 0.29 | 0.21 | 0.37 | 0.00 |
| HECO | 0.06 | 0.19 | 0.10 | 0.28 | 0.00 |
| HEDI | 0.26 | 0.54 | 0.49 | 0.59 | 0.00 |
| HEEX | 0.16 | 0.35 | 0.27 | 0.41 | 0.00 |
| HERA | 0.07 | 0.07 | 0.01 | 0.14 | 1.00 |
| HOCA | 0.29 | 0.57 | 0.52 | 0.61 | 0.00 |
| LACA | 0.15 | 0.57 | 0.52 | 0.62 | 0.00 |
| LAMI | 0.22 | 0.62 | 0.58 | 0.66 | 0.00 |
| LEMI | 0.16 | 0.61 | 0.56 | 0.65 | 0.00 |
| LIBI | 0.14 | 0.45 | 0.38 | 0.51 | 0.00 |
| LIDI | 0.12 | 0.28 | 0.20 | 0.36 | 0.00 |
| LOCI | 0.10 | 0.50 | 0.44 | 0.56 | 0.00 |
| LOHU | 0.19 | 0.60 | 0.56 | 0.64 | 0.00 |
| LOMA | 0.00 | 0.04 | 0.00 | 0.11 | 0.01 |
| LOMI | 0.12 | 0.61 | 0.56 | 0.65 | 0.00 |
| LUMI | 0.00 | 0.02 | 0.00 | 0.07 | 0.58 |
| MEIN | 0.17 | 0.37 | 0.30 | 0.44 | 0.00 |
| MICA | 0.00 | 0.02 | 0.00 | 0.07 | 0.29 |
| MICDO | 0.01 | 0.07 | 0.01 | 0.15 | 0.01 |
| MIGU | 0.35 | 0.70 | 0.66 | 0.73 | 0.00 |
| MILA | 0.18 | 0.59 | 0.55 | 0.64 | 0.00 |
| MINDO | 0.02 | 0.04 | 0.00 | 0.11 | 0.51 |
| MINU | 0.24 | 0.51 | 0.45 | 0.56 | 0.00 |
| NAJE | 0.26 | 0.60 | 0.56 | 0.64 | 0.00 |
| NEMO | 0.22 | 0.61 | 0.56 | 0.65 | 0.00 |
| PLST | 0.17 | 0.58 | 0.53 | 0.62 | 0.00 |
| RACA | 0.16 | 0.55 | 0.50 | 0.60 | 0.00 |
| SCSI | 0.21 | 0.48 | 0.43 | 0.54 | 0.00 |
| SECL | 0.11 | 0.61 | 0.56 | 0.65 | 0.00 |
| SIBE | 0.22 | 0.55 | 0.51 | 0.60 | 0.00 |
| SIDI | 0.18 | 0.60 | 0.55 | 0.64 | 0.00 |
| STAL | 0.25 | 0.62 | 0.58 | 0.66 | 0.00 |
| STBR | 0.33 | 0.63 | 0.59 | 0.67 | 0.00 |
| THMA | 0.01 | 0.04 | 0.00 | 0.09 | 0.21 |
| TRLA | 0.25 | 0.54 | 0.48 | 0.59 | 0.00 |
| TRLX | 0.20 | 0.61 | 0.56 | 0.65 | 0.00 |
| TROB | 0.17 | 0.62 | 0.57 | 0.66 | 0.00 |
| TRPE | 0.35 | 0.64 | 0.60 | 0.68 | 0.00 |
| VETCH | 0.12 | 0.07 | 0.02 | 0.14 | 0.21 |
| VIDO | 0.10 | 0.14 | 0.06 | 0.23 | 0.27 |
| ZIVE | 0.08 | 0.63 | 0.59 | 0.67 | 0.00 |

**Table S5.**
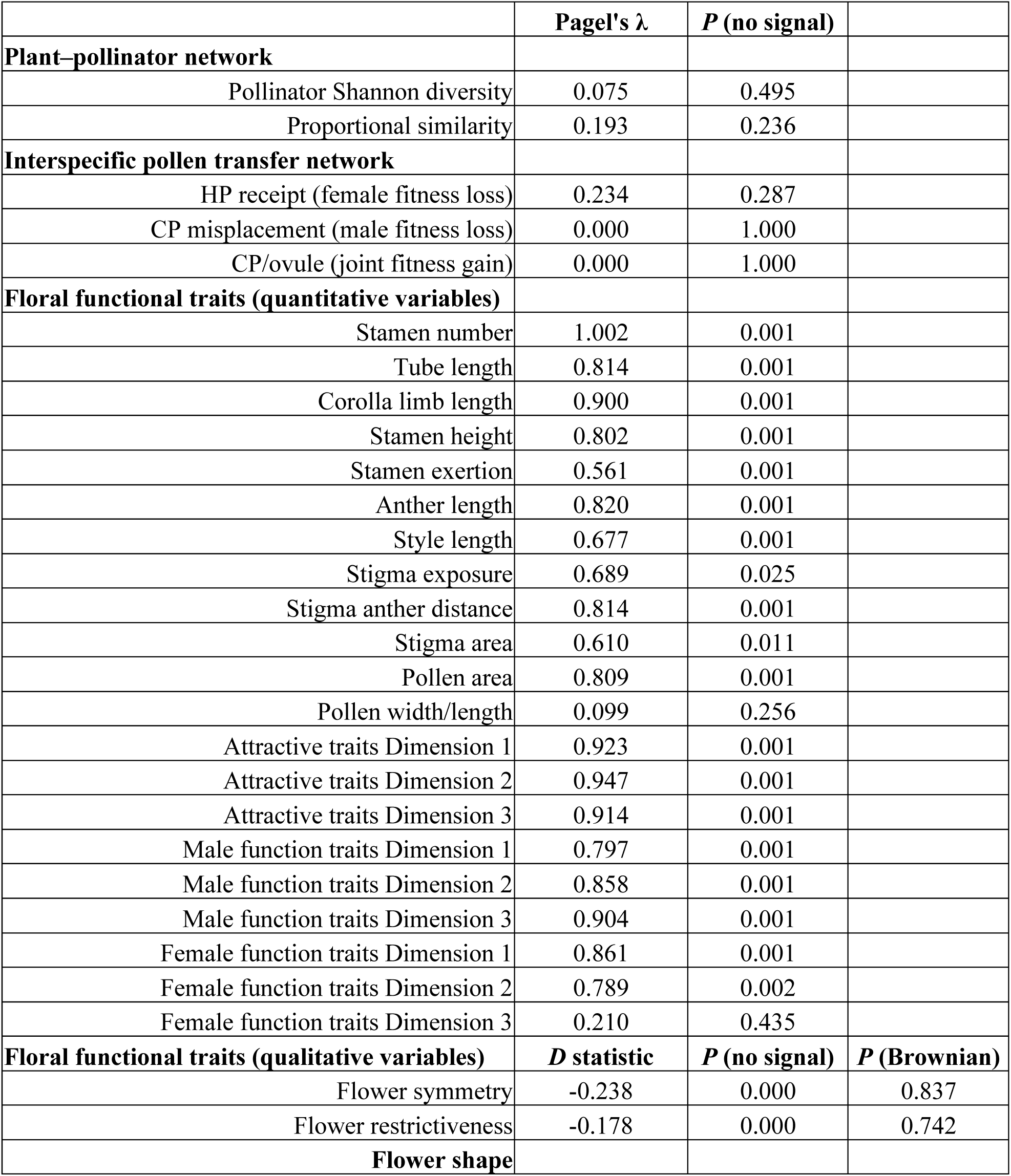
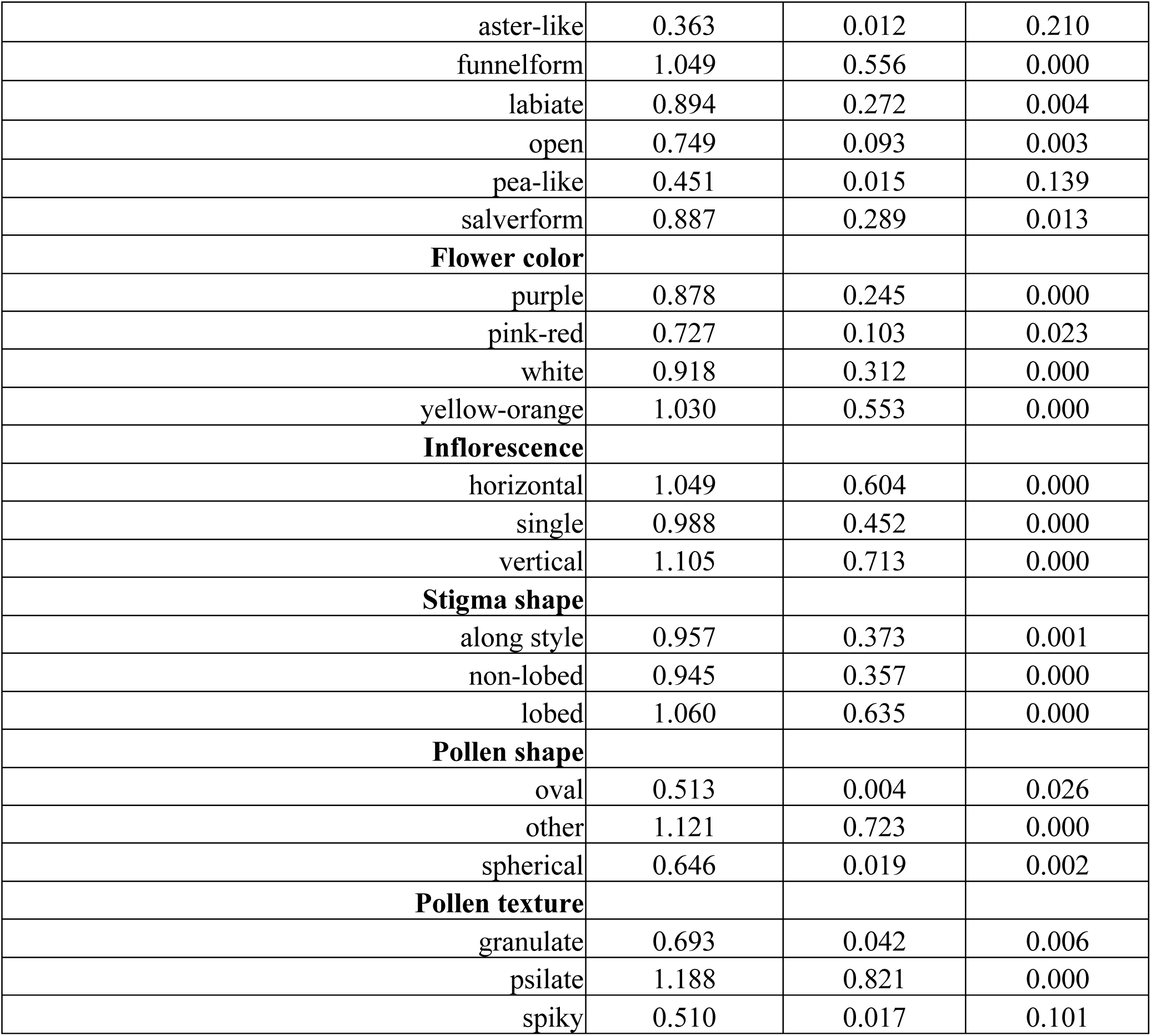
Phylogenetic signals of plant–pollinator interactions, interspecific pollen transfer network metrics and floral functional traits. Quantitative variables were examined using Pagel’s λ implemented in the R package phylosignal (*66*), and qualitative variables used the *D* statistic implemented in the package caper (*68*). The lack of a phylogenetic signal [*P*(no signal) > 0.05] is indicated when Pagel’s λ is not significantly above zero or the *D* statistic does not significantly deviate from one. When the *D* statistic does not significantly deviate from zero, it indicates Brownian motion evolution [*P*(Brownian) > 0.05]. HP, heterospecific pollen; CP, conspecific pollen.

